# Crystallographic and cryogenic electron microscopic structures and enzymatic characterization of sulfur oxygenase reductase from *Sulfurisphaera tokodaii*

**DOI:** 10.1101/2020.05.03.074773

**Authors:** Yuta Sato, Takashi Yabuki, Naruhiko Adachi, Toshio Moriya, Takatoshi Arakawa, Masato Kawasaki, Chihaya Yamada, Toshiya Senda, Shinya Fushinobu, Takayoshi Wakagi

**Affiliations:** Department of Biotechnology, The University of Tokyo, 1-1-1 Yayoi, Bunkyo-ku, Tokyo 113-8657, Japan; Structural Biology Research Center, Institute of Materials Structure Science, High Energy Accelerator Research Organization (KEK), 1-1 Oho, Tsukuba, Ibaraki, 305-0801, Japan; Collaborative Research Institute for Innovative Microbiology, The University of Tokyo, 1-1-1 Yayoi, Bunkyo-ku, Tokyo, 113-8657, Japan

**Keywords:** X-ray crystallography, cryogenic electron microscopy, sulfur metabolism, archaea, nonheme mononuclear iron center

## Abstract

Sulfur oxygenase reductases (SORs) are present in thermophilic and mesophilic archaea and bacteria, and catalyze oxygen-dependent oxygenation and disproportionation of elemental sulfur. SOR has a hollow, spherical homo-24-mer structure and reactions take place at active sites inside the chamber. The crystal structures of SORs from two *Acidianus* species have been reported. However, the states of the active site components (mononuclear iron and cysteines) and the entry and exit paths of the substrate and products are still in dispute. Here, we report the biochemical and structural characterizations of SORs from the thermoacidophilic archaeon *Sulfurisphaera tokodaii* (StSOR) and present high-resolution structures determined by X-ray crystallography and cryogenic electron microscopy (cryo-EM). The crystal structure of StSOR was determined at 1.73 Å resolution. At the catalytic center, iron is ligated to His86, His90, Glu114, and two water molecules. Three conserved cysteines in the cavity are located 9.5∼13 Å from the iron and were observed as free thiol forms. A mutational analysis indicated that the iron and one of the cysteines (Cys31) were essential for both activities and the other two cysteines (Cys101 and Cys104) had a supportive role. The cryo-EM structure was determined at 2.24 Å resolution using an instrument operating at 200 kV. The two structures determined by different methodologies showed similar main chain traces, but the maps exhibited different features at catalytically important components. Given the high resolution achieved in this study, StSOR was shown to be a good benchmark sample for cryo-EM measurements.

**Highlights:**

- Sulfur oxygenase reductase (SOR) was biochemically and structurally characterized.
- High resolution structures of SOR were determined by crystallography and cryo-EM.
- Twenty-four identical subunits of SOR form a hollow sphere.
- Catalytic components exhibited different features in the crystal and cryo-EM structures.

## 1. Introduction

In addition to geochemical activities, archaea and bacteria play significant roles in both oxidation and reduction of sulfur compounds in the global sulfur cycle. Oxidation of inorganic sulfur compounds under anaerobic conditions is the most important energy-yielding reaction for microorganisms living in those critical environments (Ghosh and Dam, 2009; Offre et al., 2013). The iron-sulfur world hypothesis postulates the emergence of the origin of life on the surfaces of inorganic sulfur, iron, and other metals under anaerobic and high temperature environments (Wächtershäuser, 2007). Thermophilic archaea and bacteria are located near the root of the phylogenetic tree, based on comparison of 16S rRNA and homologous proteins (Woese et al., 1990), and these microorganisms are thought to retain some properties of the last universal common ancestor (Weiss et al., 2016). In hydrothermal vents, solfataras, volcanic hot springs and volcanic environments all over the world, thermophilic archaea and bacteria grow under these high temperature and acidic conditions (Schönheit and Schäfer, 1995). Such thermoacidophiles belonging to the order Sulfolobales show biotechnological possibility for application to biomining, where metal sulfides in insoluble ore are recovered through the sulfur-metabolizing process of the organisms (Mangold et al., 2011; Urbieta et al., 2017).

One of the key enzymes in these processes is sulfur oxygenase reductase (SOR, EC 1.13.11.55). SORs are present in a part of thermophilic and mesophilic archaea and in bacteria such as *Acidianus ambivalens* (AaSOR) (Kletzin, 1989; Urich et al., 2004), *Acidianus tengchongensis* (AtSOR) (Chen et al., 2005; Sun et al., 2003), *Aquifex aeolicus* (AqSOR) (Pelletier et al., 2008), *Halothiobacillus neapolitanus* (HnSOR) (Veith et al., 2012), *Sulfobacillus thermosulfidooxidans* (SbSOR) (Janosch et al., 2015), and *Thioalkalivibrio paradoxus* (TpSOR) (Rühl et al., 2017). SOR catalyzes the oxygen-dependent sulfur oxygenation and disproportionation (both oxidation and reduction) reactions to produce sulfite and hydrogen sulfide and thiosulfate is concomitantly produced by a non-enzymatic reaction:

1. Oxygenation: S + O_2_ + H_2_O → HSO_3_^−^ + H^+^
2. Disproportionation: 3S + 3H_2_O → HSO_3_^−^ + 2HS^−^ + 3H^+^
3. Sum: 4S + O_2_ + 4H_2_O → 2HSO_3_^−^ + 2HS^−^ + 4H^+^
4. Non-enzymatic: S + HSO_3_^−^ → S_2_O_3_^2−^ + H^+^

In the genome of the thermoacidophilic archaeon *Sulfurisphaera tokodaii* (former name *Sulfolobus tokodaii*) strain 7 isolated from Beppu Hot Springs in Japan (Kawarabayasi et al., 2001; Tsuboi et al., 2018), a homologous *sor* gene (*st1127*) has been found. Among the characterized SORs, SOR from *S. tokodaii* (ST1127, StSOR) is most similar to the SORs from *Acidianus* species (amino acid sequence identity with AaSOR = 68.6%, Fig. 1). A Blast search based on the deduced amino acid sequence showed that highly similar proteins with more than 50% identity are present in other members of Sulfolobaceae such as *Sulfurisphaera ohwakuensis, Sulfuracidifex metallicus* (former name *Sulfolobus metallicus*), *A. ambivalens, Acidianus brierleyi, Acidianus hospitalis, Acidianus infernus*, and *Acidianus sulfidivorans*. In contrast, *sor*-like genes are not present in some Sulfolobales members such as *Saccharolobus solfataricus* (former name *Sulfolobus solfataricus*) P2 and *Sulfolobus acidocaldarius* strain DSM639.

**Fig. 1.**
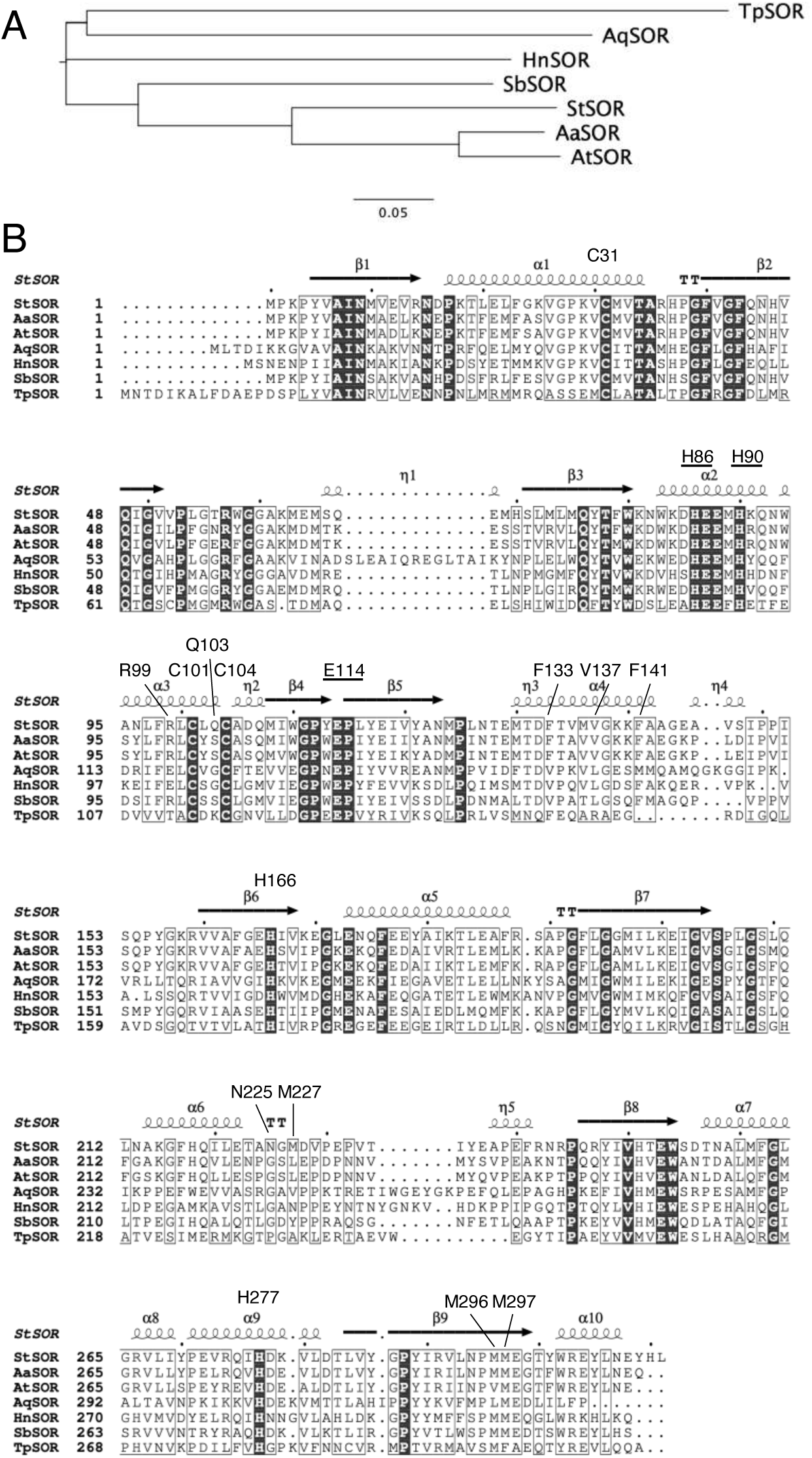
A phylogenetic tree (A) and amino acid sequence alignment (B) of StSOR and characterized SORs. AaSOR, *Acidianus ambivalens* SOR; AtSOR, *Acidianus tengchongensis* SOR; AqSOR, *Aquifex aeolicus* SOR; HnSOR, *Halothiobacillus neapolitanus* SOR; SbSOR, *Sulfobacillus thermosulfidooxidans* SOR; and TpSOR, *Thioalkalivibrio paradoxus* SOR. (B) Secondary structures of StSOR are shown above the sequences. Iron ligands (underlined) and other residues referred in the text are also indicated.

Elemental sulfur (S^0^) can support the growth of *S. solfataricus* P2, *S. acidocaldarius* DSM639 (Grogan, 1989), and *S. ohwakuensis* strain TA-1 (Kurosawa et al., 1998). *S. tokodaii* strain 7 is an obligate aerobe and grows optimally at 80°C and pH 2.5–3 under aerobic and chemoheterotrophic conditions but autotrophic growth on S^0^ has not been reported so far (Suzuki et al., 2002). This organism is applied to oxidize H_2_S to H_2_SO_4_ at a geothermal power station in Kyusyu, Japan (Takeuchi and Hirowatari, 1999).

SOR forms a homo-oligomeric complex. Each protomer contains approximately 300 amino acids and one non-heme mononuclear iron center (Kletzin, 1992; Urich et al., 2004). The crystal structures of AaSOR and AtSOR have revealed that the sphere-shaped tetracosamer (24-mer) is organized with *432* symmetry (Li et al., 2008; Urich et al., 2006). The molecular organization is consistent with the previous electron micrographic observation for a negatively stained AaSOR, in which the oligomer was seen as particles of 15.5 nm diameter (Urich et al., 2004).

For further elucidation of the relationship between SOR structure and function, we report here the enzymatic characterizations of StSOR and its three-dimensional structures determined by X-ray crystallography and cryogenic electron microscopy (cryo-EM), as well as functional analyses of variant enzymes using site-directed mutagenesis. Our study demonstrated in vitro functionality of StSOR and several structural differences at the active site in comparison with SORs from *Acidianus* species. We also found several differences in the X-ray crystallographic and cryo-EM maps at the components involved in the enzymatic function.

## 2. Materials and methods

### 2.1 Protein preparation

The open reading frame of *st1127* was amplified by PCR using *S. tokodaii* genomic DNA as a template with *PfuTurbo* DNA polymerase (Agilent Technologies, Santa Clara, CA, USA). The following primers were used: 5’-GTAATGACCA<u>CATATG</u>CCGAAACCATACGTAGCAATAAAC-3’ (forward) and 5’-CATAGTTCTA<u>CTCGAG</u>GGTTATAAATGATATTC-3’ (reverse). The NdeI and EcoRI restriction sites are underlined. The PCR product was digested with restriction enzymes and then ligated into pET-17b (Merck Millipore, Burlington, MA, USA) to construct the expression vector pStSOR. The sequence of the insert was confirmed by DNA sequencing.

For enzymatic and biochemical characterizations, the recombinant enzyme of StSOR was expressed and purified as follows. *E. coli* BL21 (DE3) pLysS (Agilent Technologies) cells harboring pStSOR were grown in 100 mL of Luria-Bertani medium containing 100 μg mL^−1^ ampicillin and 30 μg mL^−1^ chloramphenicol overnight at 37°C. The cell culture (80 mL) was inoculated into 8 liters of the same medium and then grown for 9 h at 37°C without induction. Expressed cells were harvested by centrifugation and suspended in 10 mM Tris-HCl (pH 8.6). The cells were disrupted by sonication and the supernatant after centrifugation was heated at 75°C for 15 min. After centrifugation, the supernatant was purified to homogeneity by column chromatography employing Q-Sepharose Fast Flow (GE Healthcare Life Sciences, Buckinghamshire, UK), Hydroxyapatite (Bio-Rad Laboratories, Hercules, CA, USA), Mono Q 10/10 (GE Healthcare Life Sciences), and HiLoad 16/600 Superdex 200 pg (GE Healthcare Life Sciences) sequentially. The molecular weight standards used for gel filtration chromatography were thyroglobulin (669 k), pyruvate kinase (237 k), ovalbumin (44 k), and RNaseA (14 k).

For structural analyses, the recombinant enzyme was expressed and purified as follows. *E. coli* BL21-AI (Thermo Fisher Scientific, Waltham, MA, USA) cells harboring pStSOR were grown at 37°C in 3 liters of Luria-Bertani medium containing 20 μg mL^−1^ ampicillin until the absorbance at 600 nm reached 0.6. Protein expression was induced by adding 0.1 mM isopropyl-β-D-thiogalactopyranoside and 0.5% (w/v) L-arabinose to the medium. The culture was harvested after an additional incubation for 24 h at 37°C. After cell disruption and heat treatment at 70°C for 1 h and centrifugation, the supernatant was purified by gel filtration using a HiLoad 16/600 Superdex 200 pg column. The sample was used for cryo-EM analysis at this step. For crystallization, the protein was further purified using a Mono Q 16/10 column. The protein concentrations were determined by a BCA protein assay kit (Thermo Fischer Scientific) with bovine serum albumin as the standard. The protein concentration measurements were based on the absorbance at 280 nm and a theoretical extinction coefficient calculated from the amino acid sequence (60,850 M^−1^ cm^−1^) was also used for purified protein samples to check the consistency.

### 2.2 Enzyme assay and quantification of free thiols

The SOR activity was measured by generally following the methods presented by Kletzin et al. (Kletzin, 1989; Urich et al., 2004). The enzymatic reaction was performed aerobically at 80°C in round-bottom stoppered test tubes reciprocally shaken at 120 rpm. The reaction was started by adding an enzyme solution (50 μL) to the assay solution (2,950 μL) containing 2% (w/v) sulfur, 0.005% Tween 20, and 20 mM MES-NaOH (pH 6.0). S^0^ was suspended and dispersed with an ultrasonic homogenizer before the reaction. The reaction was stopped by cooling the sample on ice. Fractions (1 mL) of the reaction mixture were recovered in test tubes and the supernatant was used for the product analysis after centrifugation at 2,300 *g* for 2 min at 4°C. One unit of activity was defined as μmoles of sulfite plus thiosulfate (oxygenase) or of hydrogen sulfide (reductase) formed per min.

Sulfite concentrations were determined by the previously described method (Kletzin, 1989; Suzuki and Silver, 1966). The assay solution (350 μL) contained 35 μL of 0.3 M H_2_SO_4_, 10 μL of 0.08% (w/v) Fuchsin solution (Sigma-Aldrich Co., St. Louis, MO, USA), 5 μL of 1.5% (v/v) formaldehyde, 150 μL of double-distilled water, and 150 μL of the sample. The absorbance at 570 nm was measured after the coloring reaction. The thiosulfate concentration was determined by the previously described method (Impert et al., 2003; Kletzin, 1989). The assay solution (450 μL) contained 350 μL of coloring reagent (1:1 mixture of 0.0024% (w/v) methylene blue and concentrated HCl) and 100 μL of the sample. After incubation in a sealed tube for 2 h, the absorbance at 666 nm was measured. The molar extinction constant of methylene blue (7.8 ×10^4^ M^−1^cm^−1^) was used (Hay et al., 1981). The hydrogen sulfide concentration was determined by the previously described method (King and Morris, 1967; Kletzin, 1989). The assay solution (580 μL) contained 300 μL of 0.1% (w/v) *N,N*-dimethyl-*p*-phenylenediamine in 5.5 N HCl, 80 μL of 23 mM FeCl_3_ in 1.2 N HCl, and 200 μL of the sample. The formation of methylene blue was measured by the absorbance at 666 nm.

Enzymatic thermal stability was evaluated by measuring the activities under standard assay conditions after incubation of the enzyme (0.5 mg/mL) at 25–100°C for 30 min in a 10 mM Na-phosphate buffer (pH 8.0). The effects of temperature and pH on the activities were evaluated by measuring the activities under standard assay conditions by changing the temperature between 40 and 100°C or by substituting the buffer with the following 20 mM buffers: citrate-HCl (pH 4.0–6.0), MES-NaOH (pH 5.5–7.0), and Tris-HCl (pH 7.5– 9.0). To investigate the effects of additives (inhibitors or denaturants), the protein sample was incubated with each concentration of additive at 25°C for 15 min before use in the standard assays and the remaining activity was compared with the sample without additives. Free thiols in the StSOR protein were quantified using 5,5’-dithiobis(2-nitrobenzoic acid) (DTNB) or *p*-chloromercuribenzoate (*p*CMB) by the previously described method (Shoun, 1990).

### 2.3 Spectroscopy

UV-visible absorption spectra were collected with a JASCO V-560 spectrophotometer. Circular dichroism (CD) spectra were recorded with a JASCO J720 spectrophotometer using a cylindrical cell with a 1 mm light path. The contents of secondary protein structure were estimated by the K2D3 server (Louis-Jeune et al., 2012). Inductively coupled plasma atomic emission spectroscopy (ICPAES) was measured with an SPS1200VR (SEIKO Instruments, Chiba, Japan). Before the ICPAES measurements, the protein sample was dialyzed against double-distilled water, dried by centrifugation under reduced pressure, dissolved in 60 μL of 61% HNO_3_, and then heated at 90°C for 1 h. After drying, it was dissolved with 2.5 mL of 0.1 N HNO_3_. The amount of iron was calculated using appropriate standard solutions.

### 2.4 Site-directed mutants

Mutants of StSOR were constructed with a QuikChange site-directed mutagenesis kit (Agilent Technologies) using the following and complementary oligonucleotides: 5’-GACCTAAAGTA<u>GC</u>CATGGTCACAGC-3’ (C31A), 5’-GAATTGGAAAGAT<u>GC</u>TGAGGAAATGC-3’ (H86A), 5’-GAGGAAATG<u>GC</u>TAAACAGAACTGG-3’ (H90A), 5’-CTTTAGGTTA<u>GC</u>TCTACAATGTGC-3’ (C101A), 5’-GTTATGTCTACAA<u>GC</u>TGCAGACCAAATG-3’ (C104A), and 5’-GGACCTTATG<u>C</u>ACCTTTATATGAAATAG-3’ (E114A). The mutations were confirmed by DNA sequencing. The mutant enzymes were expressed and purified with heat treatment and column chromatography steps using Q-Sepharose and HiLoad 16/600 Superdex 200 pg columns.

### 2.5 Crystallography

StSOR crystals were grown at 20°C using the sitting drop vapor diffusion method by mixing 1.0 μL of 10 mg/mL protein solution in 10 mM Tris-HCl (pH 8.0) with an equal volume of reservoir solution containing 1.2 M ammonium sulfate, 100 mM MES-NaOH (pH 6.1), and 4% (w/v) dioxane. The crystals were cryoprotected by the reservoir solution supplemented with 25% (w/v) glycerol and were flash cooled by dipping into liquid nitrogen. The X-ray diffraction dataset was collected at a wavelength of 1.0 Å at 100 K on beamline BL-5A at the Photon Factory of the High Energy Accelerator Research Organization (KEK, Tsukuba, Japan). The initial phase was determined by the molecular replacement method using Molrep (Vagin and Teplyakov, 2010) and the crystal structure of AaSOR (PDB: 2CB2) was used as the search model. Manual model building was performed using Coot (Emsley et al., 2010). Crystallographic refinement was carried out using Refmac5 (Murshudov et al., 2011) with local non-crystallographic symmetry restraints and Babinet’s bulk solvent type scaling was applied. Molprobity-based “Comprehensive validation” (Williams et al., 2018) in PHENIX was used for structural validation. A Polder map (Liebschner et al., 2017) was created using PHENIX. Molecular graphic images were prepared using PyMOL (Schrödinger, LLC, New York, USA).

### 2.6 Cryo-EM

For cryo-grid preparation, 3 µL of the sample with 10 mg/mL concentration was dissolved in 20 mM Tris-HCl (pH 8.0) and 150 mM NaCl. The sample was applied onto a holey carbon grid (Quantifoil, Cu, R1.2/1.3, 300 mesh) which was rendered hydrophilic by a 30 s glow-discharge in air (11 mA current) with PIB-10 (Vacuum Device Inc., Ibaraki, Japan). The grid was blotted for 5 s (blot force 20) at 18°C and 100% humidity and was flash-frozen in liquid ethane using Vitrobot Mark IV (Thermo Fisher Scientific). For automated data collection, 2,558 micrographs were acquired on a Talos Arctica (Thermo Fisher Scientific) microscope operating at 200 kV in nanoprobe mode using EPU software. The movie micrographs were collected on a 4k x 4k using a Falcon 3EC direct electron detector (electron counting mode) at a nominal magnification of 150,000 (0.69 Å/pixel). Fifty movie fractions were recorded at a dose of 1.00 electrons per Å^2^ per fraction corresponding to a total dose of 50 e^−^/Å^2^. The defocus steps used were –0.3, –0.6, –0.9, – 1.2, and –1.5 µm.

For data processing, the movie fractions were aligned, dose-weighted, and averaged using Motioncor2 (S. Q. Zheng et al., 2017). The micrographs whose total accumulated motion was larger than 100 Å were discarded. The non-weighted movie sums were used for Contrast Transfer Function (CTF) estimation with CTFFIND4 (Rohou and Grigorieff, 2015) and the Gctf program (Zhang, 2016). The images whose CTF max resolution was better than 5.5 Å were selected. In addition, micrographs showing obvious indications of ice crystallization were manually discarded by visual inspection. The particles were picked fully automatically from 2,387 micrographs using SPHIRE-crYOLO (Moriya et al., 2017; Wagner et al., 2019). The subsequent processes, namely, 2D classification, *ab initio* reconstruction, 3D classification, 3D refinement, CTF refinement, and Bayesian polishing were performed by using RELION-3 (Zivanov et al., 2018). A stack of 305,182 particle images was extracted from the dose-weighted sum micrographs while rescaling to 2.76 Å/pixel with a 96-pixel box size and were subjected to two consecutive runs of 2D classification. Next, 160,571 particles, which corresponded to the best classes of the 2nd run, were selected for *ab initio* reconstruction (asymmetry and single expected class) and 176,437 particles were selected with more relaxed criteria for the subsequent 3D classification (2 expected classes). The generated *ab initio* map was imposed octahedral symmetry and used as an initial model for the 3D classification. The 3D volume and 152,484 particles of the best 3D class were used for the subsequent 3D refinements with octahedral symmetry. The refined volume and particle images were rescaled to 0.69 Å/pixel with a 400-pixel box size and were 3D auto-refined (octahedral symmetry) twice (the 1st run was without and the 2nd run was with a soft-edged 3D mask). Then, 152,457 selected particles were again re-centered and re-extracted without changing the rescaling settings. The cycle of CTF refinement and Bayesian polishing was repeated four times. 3D refinement with octahedral symmetry was executed after each CTF refinement and Bayesian polishing step. Then, the pixel size was checked by comparing the cryo-EM structure with the atomic-coordinate model of the crystal structure using the “Fit in Map” tool in UCSF Chimera (Pettersen et al., 2004) and was calibrated to 0.676 Å/pixel. After the calibration, 146,465 selected particles were again re-centered and re-extracted with a 480-pixel box size and were subjected to subsequent 3D refinement (octahedral symmetry) and the two cycles of CTF refinement and Bayesian polishing steps. Then, no-alignment 3D classification was conducted (octahedral symmetry, 2 expected classes, T = 16), and 85,621 particles were selected by choosing the best 3D class. The last 3D refinement (octahedral symmetry) generated the final result at 2.24 Å resolution. The gold-standard FSC resolution with 0.143 criterion (Rosenthal and Henderson, 2003) was used as the global resolution estimation. The local resolution was estimated using RELION-3’s own implementation of ResMap (Kucukelbir et al., 2014). UCSF Chimera and e2display.py of EMAN2 (Tang et al., 2007) were used for the visualization.

For initial model reconstruction, “Map_to_Model” (Terwilliger et al., 2018) in PHENIX was used. During the procedure, the model was refined with the amino acid sequence and octahedral symmetry of StSOR, while the crystal structure was not referenced to avoid model bias. The reconstructed model was further refined both manually on Coot and automatically with “Real-Space Refine” (Afonine et al., 2018) in PHENIX. Validation of the refined model was carried out using “Comprehensive validation” in PHENIX. UCSF Chimera and Coot were used for visualization.

## 3. Results

### 3.1 Protein purification and characterization

For biochemical and enzymatic characterizations, the recombinant enzyme of the intact StSOR polypeptide (ST1127, 311 amino acids) without any tag sequence was expressed in *E. coli* and purified to homogeneity by heat treatment and chromatography steps with four different columns (Fig. S1A). On sodium dodecyl sulfate (SDS)-denatured polyacrylamide gel electrophoresis (PAGE), a major single band appeared at a position of apparent molecular mass of 38 kDa, which was comparable to the predicted molecular mass of the recombinant protein (35.7 kDa). At the gel filtration step, the protein exhibited a peak corresponding to a molecular mass of 539 kDa. This value corresponds to 15-to 16-mer and was not significantly different from the reported molecular masses of SORs from other organisms estimated using the same method (∼550 kDa) (Emmel et al., 1986; Kletzin, 1989; Sun et al., 2003; Urich et al., 2004). These results suggest that the oligomeric numbers of SORs estimated by size exclusion chromatography (hydrodynamic volume) are usually underestimated due to their spherical shapes.

### 3.2 Enzymatic properties

The oxygenase and reductase activities of StSOR were measured by the production of sulfite (HSO_3_^−^) plus thiosulfate (S_2_O_3_^2−^) and that of hydrogen sulfide (H_2_S), respectively. The purified recombinant protein sample exhibited specific activities of 10.5 U/mg oxygenase and 0.95 U/mg reductase, respectively (Table 1). A purified recombinant protein sample of AaSOR was reported to have specific activities of 2.96 U/mg oxygenase and 0.60 U/mg reductase (Urich et al., 2004). The remaining StSOR activities after 30 min of incubation at varying temperatures were measured (Fig. 2A). The oxygenase activity gradually decreased while the reductase activity remained relatively stable. After 30 min of incubation at 80°C, the remaining oxygenase and reductase activities were approximately 40% and 90%, respectively and suggest that the oxygenation reaction of StSOR is less thermally tolerant than the disproportionation reaction. The apparent optimum StSOR temperature was 80°C (Fig. 2B). The optimum temperatures of SORs from other thermophiles were reported to be 70–85°C (Kletzin, 1989; Pelletier et al., 2008; Rühl et al., 2017; Sun et al., 2003; Veith et al., 2012). The optimum pH of StSOR was 6.0 for both oxygenase and reductase activities, and oxygenase activity significantly dropped at pH levels above 6.5 (Fig. 2C). StSOR is suggested to function optimally in *S. tokodaii* cells because the intracellular pH of another thermoacidophilic archaeon, *S. acidocaldarius*, is estimated to be approximately 6.5 (Moll and Schäfer, 1988). The optimum pH values for AaSOR and AtSOR are ∼7 and 5.0, respectively (Kletzin, 1989; Sun et al., 2003). A similar activity profile with a significant reduction at neutral and alkaline pH values (above 6.0) was also observed for AtSOR (Sun et al., 2003).

**Table 1.** Enzymatic activities of StSOR and its mutants.

| Enzyme | Oxygenase (U/mg) | Reductase (U/mg) |
| --- | --- | --- |
| Wild type | 10.5 | 0.95 |
| C31A | ND | ND |
| C101A | 0.93 | 0.11 |
| C104A | 0.86 | 0.14 |
| H86A | ND | ND |
| H90A | ND | ND |
| E114A | ND | ND |

Oxygenase (production of sulfite and thiosulfate) and reductase (production of hydrogen sulfide) activities were measured. ND: not detected.

**Fig. 2.**
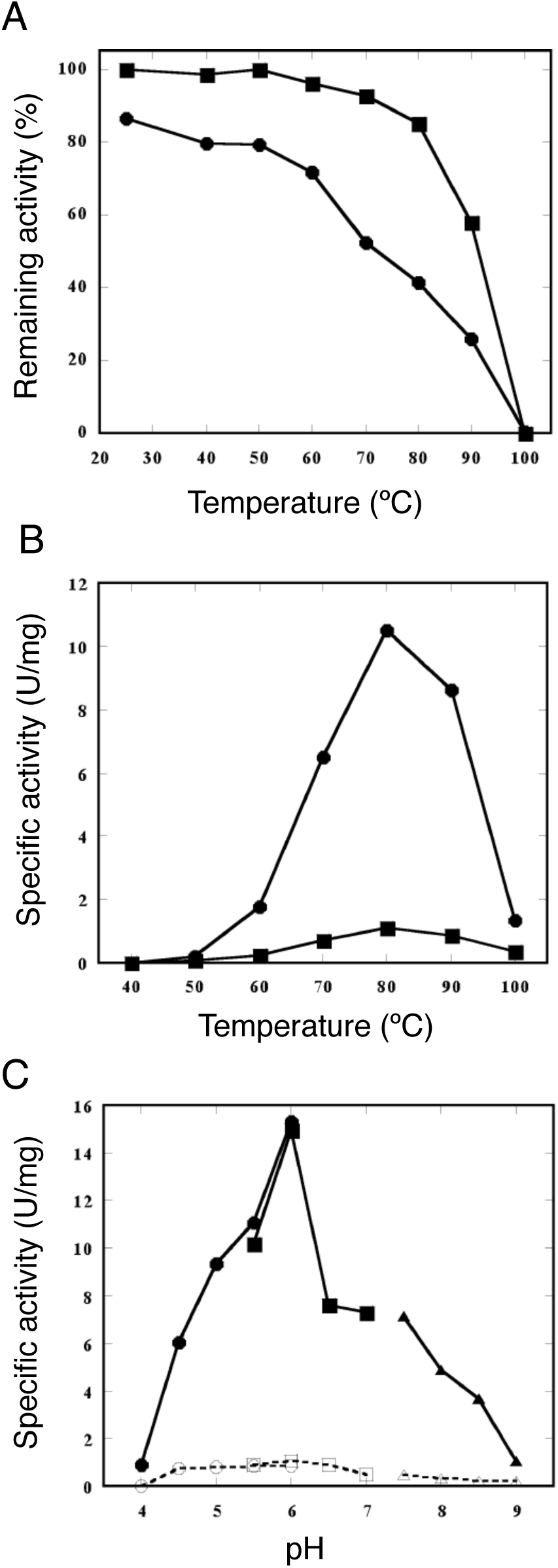
Enzymatic characterization of StSOR. (A) Thermal stability. Remaining activities after incubation at each temperature for 30 min were measured. Symbols: filled circle, oxygenase; filled square, reductase. (B) Temperature dependence measured at pH 6.0. Symbols: filled circle, oxygenase; filled square, reductase. (C) pH dependence measured at 80 °C. Symbols: filled symbols, oxygenase; open symbols, reductase; circles, citrate-HCl buffer; squares, MES-NaOH buffer; triangles, Tris-HCl buffer. Details of the assay conditions are described in the main text.

The effects of inhibitors and metal ions on the activities were measured (Fig. S2A). Sodium azide (NaN_3_) did not affect the activities while potassium cyanide (KCN) moderately inhibited oxygenase activity (71% with 0.1 mM KCN). Hg^2+^ and Cu^2+^ significantly inhibited both activities, whereas Zn^2+^ significantly activated reductase activity. Reductase activity was not significantly affected or was slightly activated by Fe^2+^ and Fe^3+^ ions. On the other hand, oxygenase activity was significantly inhibited by both Fe ions. Reducing agents (dithiothreitol and *N*-ethylmaleimide) and thiol modifiers (*p*CMB and iodoacetic acid) inhibited both activities. A metal chelator, ethylenediaminetetraacetic acid, inhibited reductase activity by approximately 50%. These effects of additives on the StSOR activities were mostly similar to those for AaSOR and AtSOR (Chen et al., 2005; Kletzin, 1989). The effects of denaturants were also examined (Fig. S2B). After 30 min of incubation at 4°C, the remaining activity decreased to less than 50% in the presence of > 1 M guanidium chloride and > 2 M urea. SDS exhibited a high denaturing effect as the activity decreased significantly at concentrations of approximately 0.05 ∼ 0.5%.

In the UV-visible absorption spectra, StSOR exhibited a slight shoulder approximately 350-400 nm at high protein concentrations (Fig. S3A). The contents of the secondary structural elements estimated by a CD spectrum (Fig. S3B) were 34.93% α helix and 20.1% β strand. ICPAES measurements indicated that the purified StSOR protein sample contained 0.8 mol iron atom per monomer. The number of free thiols in the purified StSOR protein sample was estimated to be 2.8 and 2.5 mol per monomer, respectively, by the DTNB and *p*CMB methods.

### 3.3 Crystal structure and site-directed mutant analysis

A protein sample of StSOR purified by heat treatment and two successive column chromatographies (gel filtration and anion exchange) was used for crystallization (Fig. S1B, lane 3). The crystal structure was determined at 1.73 Å resolution using a crystal belonging to the cubic *I*32 space group (Table 2). The crystal contains 8 monomers per asymmetric unit and the 24-mer with a pseudo-*432* point group was generated by applying the crystallographic 3-fold rotation axis (Fig. 3A). The final model contains residues from Pro2 to Leu311 of all chains without a disordered region. The 8 chains in the asymmetric unit had virtually identical structures, as the average root mean square deviation (RMSD) and their standard deviation (SD) for the Cα atoms between all chain pairs were 0.081 ± 0.031 Å (maximum = 0.152 Å in 28 pairs). The quaternary structure of the StSOR holoenzyme (calculated molecular mass = 857.4 kDa) forms a hollow sphere. The overall shape resembles those of AaSOR and AtSOR (Li et al., 2008; Urich et al., 2006). A striking structural feature of SOR, called “chimneys”, is present in StSOR as six protrusions. Each monomer has an 8-stranded β barrel core surrounded by 10 α helices (Fig. 3B). A helix-loop region (residues 125–152) and the C-terminal helix (residues 300-311) are located above the β barrel. One chimney is formed by the association of four helix-loop regions that are arranged along a 4-fold axis. The monomer structure of StSOR was also similar to those of SORs from *Acidianus* species. The average RMSD of Cα atoms and its SD between all chain pairs of StSOR (8 chains) and AaSOR (6 chains) was 0.825 ± 0.017 Å (maximum = 0.863 Å in 48 pairs).

**Table 2.**
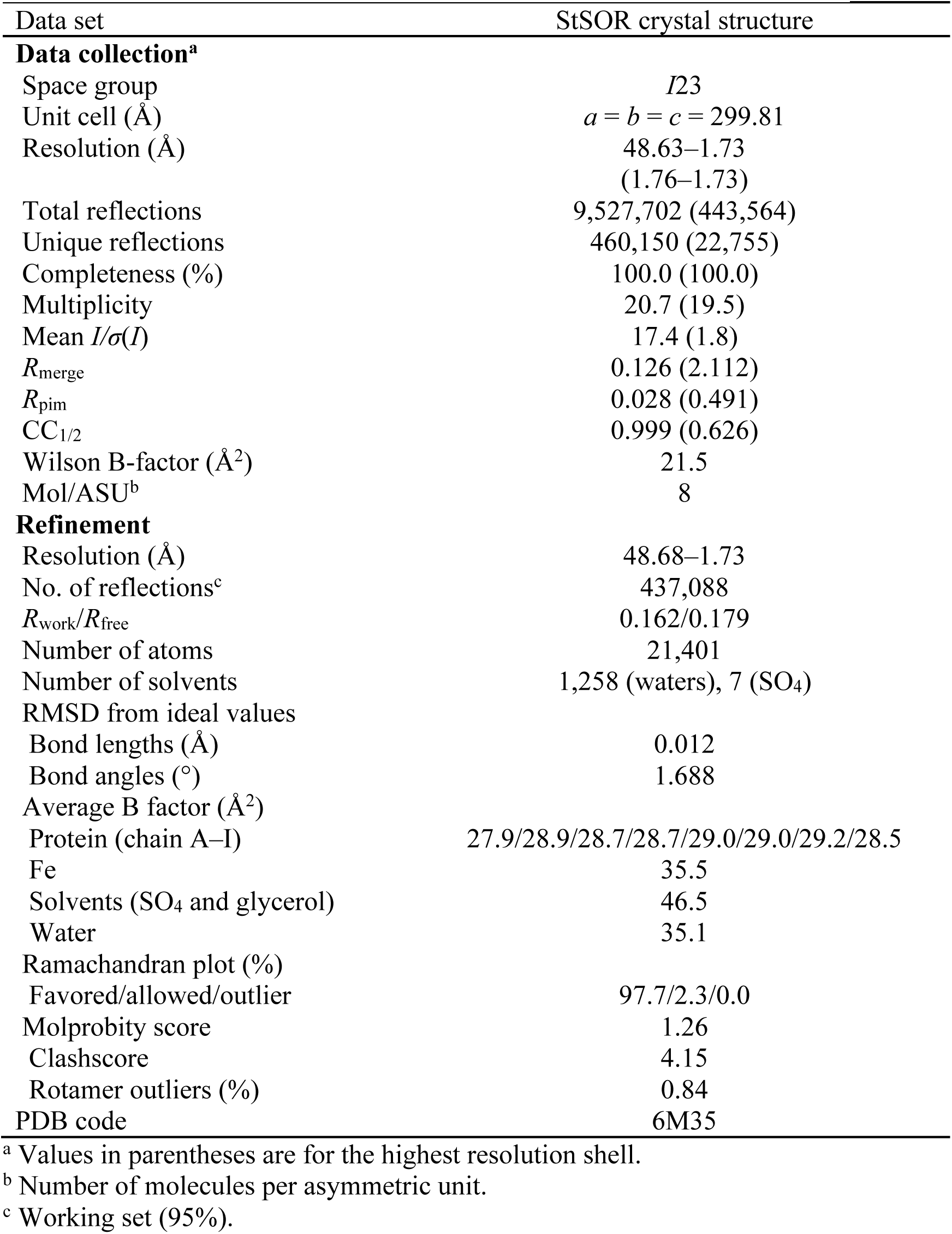
Data collection and refinement statistics for X-ray crystallography.

**Fig. 3.**
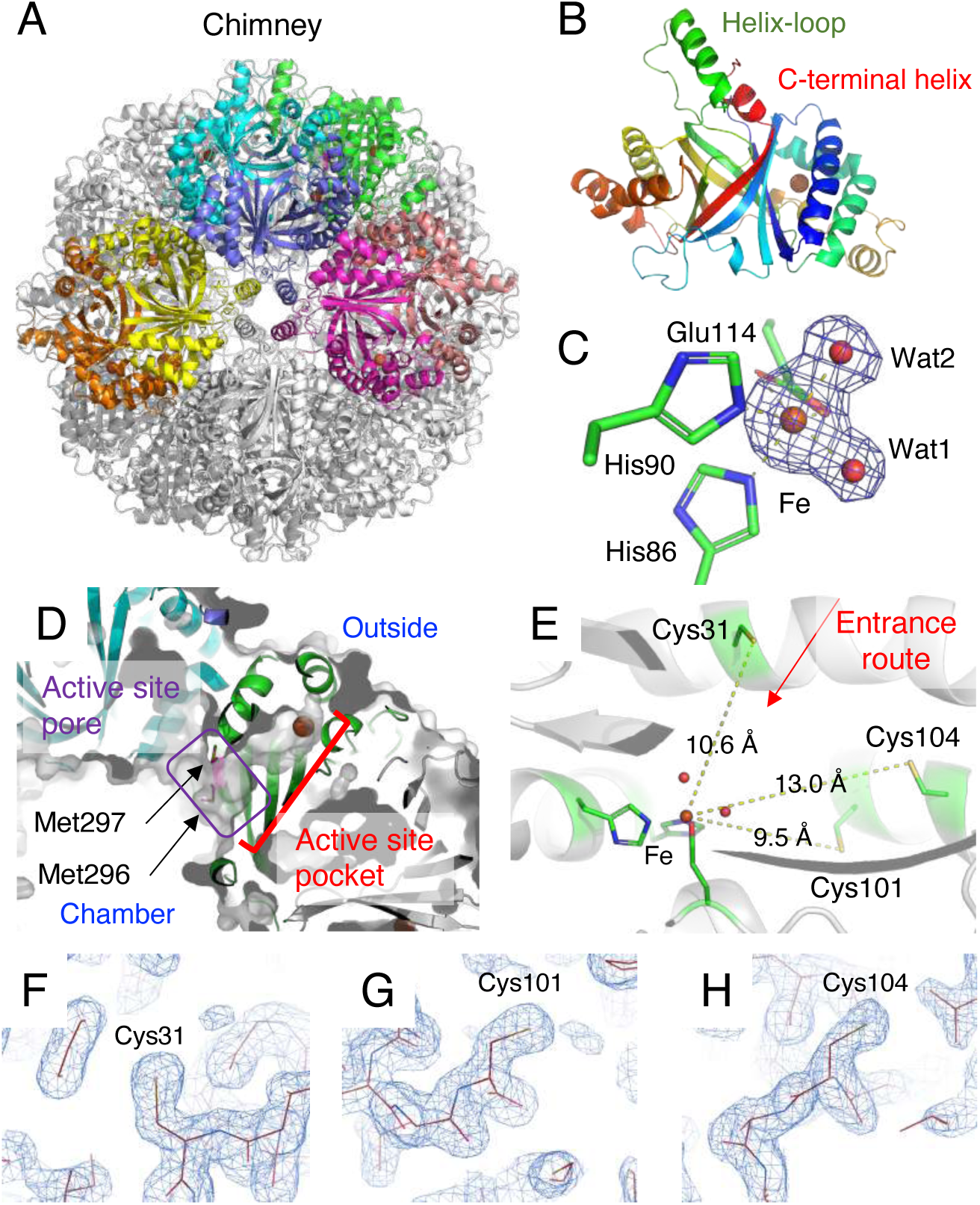
Crystal structure of StSOR. Iron atoms are shown as orange spheres. (A) Homo 24-mer structure. Eight chains in the asymmetric unit of the crystal are colored. (B) Structure of a protomer. C-terminal helix and a helix-loop region (residues 125-152), which forms the chimney, are indicated. (C) Iron center. Polder map (4.5σ) of the iron and two water atoms is shown as blue mesh. (D) The catalytic “active site pocket” containing the iron atom and the “active site pore” that connects it from the chamber are shown. (E) Cysteine residues in the active site pocket. A red arrow indicates the entrance route from the central cavity of the chamber. (F-H) Electron density map of Cys31 (F), Cys101 (G), and Cys104 (H) in the cavity. 2*F*_o_–*F*_c_ maps are shown with a contour level of 1.3σ.

The iron atom in the active site is ligated to His86, His90, Glu114 (bidentate) and two waters designated as Wat1 and Wat2 (Fig. 3C). The octahedral iron site in the StSOR crystal structure is similar to that of AaSOR (Urich et al., 2006) while the iron site of AtSOR was five-coordinated with only one water (Wat2 as the opposite ligand to His86) (Li et al., 2008). Electron paramagnetic resonance spectroscopy on AaSOR indicated that this mononuclear non-heme metal is a high-spin ferric iron (Fe^3+^) (Urich et al., 2004). The coordination distance and crystallographic temperature factors of the iron-coordinating atoms are shown in Table 3. The CheckMyMetal server indicated that the metal binding site is a distorted octahedral site of an iron and the average geometrical RMSD and SD from 8 chains is (11.0 ± 1.8)° (H. Zheng et al., 2017). The average and SD values of the bond valence and nVECSUM parameter, which indicate symmetry and completeness of the coordination sphere, of the sites in the 8 chains are 1.78 ± 0.05 and 0.080 ± 0.009, respectively. Comparison of the coordination distances with the Cambridge Structural Database suggested that the lengths are typical for Fe–N bonds and are atypical for Fe–O bonds (Fig. S4A).

**Table 3.** Coordination distances and crystallographic temperature factors of iron-coordinating atoms.

| Atom | Crystal structure<sup>a</sup> |  | Cryo-EM structure |
| --- | --- | --- | --- |
|  | Distance (Å) | B factor (Å<sup>2</sup>) | Distance (Å) |
| Fe | — | 35.53 ± 0.15 | — |
| H86 Nε2 | 2.17 ± 0.02 | 23.94 ± 1.14 | 2.22 |
| H90 Nε2 | 2.19 ± 0.02 | 32.78 ± 1.94 | 2.36 |
| E114 Oε1 | 2.29 ± 0.02 | 27.43 ± 1.05 | 2.44 |
| E114 Oε2 | 2.18 ± 0.02 | 25.20 ± 0.91 | 2.44 |
| Wat1 | 2.17 ± 0.02 | 31.71 ± 1.30 | 2.37 |
| Wat2 | 2.24 ± 0.02 | 30.82 ± 2.53 | —<sup>b</sup> |

<sup>a</sup> Average and standard deviation of the values of the 8 chains in the asymmetric unit are shown.
<sup>b</sup> Wat2 was not observed in the cryo-EM structure.

The catalysis of SOR takes place inside the hollow chamber that is separated from the cytoplasmic environment. Specifically, the active site pocket containing the iron atom is connected to the central cavity of the chamber by a narrow pore and the path is referred to as the “active site pore” (Fig. 3D) (Veith et al., 2011). In agreement with the inhibition of StSOR by cysteine modifiers (Fig. S2A), previous studies on SORs have suggested that cysteine residues near the metal center are involved in catalysis (Chen et al., 2005; Urich et al., 2005). In particular, among three cysteines in the active site pocket (Cys31, Cys101, and Cys104), Cys31 was observed as cysteine persulfide in the AaSOR crystal structure (Urich et al., 2006). StSOR also has the three conserved cysteines with distances to the iron atom of 9.5∼13.0 Å, and Cys31 is located on the route from the central cavity of the chamber to the catalytic iron (Fig. 3E). Electron density maps around the Sγ atoms of the cysteine residues clearly indicate that all are mainly present as free thiols in the StSOR crystal (Fig. 3F-H). This observation is consistent with the quantification of free thiols (2.5 ∼ 2.8 per monomer) described above and the absence of cysteine persulfide in the AtSOR crystal structure (Li et al., 2008).

To investigate the contributions of the iron ligands and cysteine residues to the catalysis of StSOR, we prepared alanine-substituted proteins of His86, His90, Glu114, Cys31, Cys101, and Cys104 by site-directed mutagenesis. As expected, both the oxygenase and reductase activities were not detected for the mutants of the iron ligands (H86A, H90A, and E114A) (Table 1). C31A also did not exhibit such activities, while C101A and C104A showed approximately 10-fold reductions relative to the wild type for both oxygenase and reductase activities. Similar effects on mutation were reported for AaSOR and AtSOR (Chen et al., 2005; Urich et al., 2005). Cys31 was suggested to play a primary role in catalysis by forming a covalent bond with the substrate, S^0^ (Urich et al., 2006; Veith et al., 2011).

### 3.4 Cryo-EM structure

The StSOR structure was determined by an independent method (cryo-EM) to further examine the active site structure. A protein sample which was partially purified by heat treatment and a gel filtration column was successfully used for cryo-EM analysis (Fig. S1B, lane 2). Table 4 summarizes the statistics of the data collection, image processing, and 3D reconstruction steps. The number of acquired micrographs was 2,558. The StSOR particles in micrographs after motion collection were easy to recognize because of the high image contrast of the 200 kV imaging (Fig. 4A). The 2D class average images reconstructed by two consecutive runs of 2D classification starting from the 305,182 particle images clearly showed the secondary-structural elements and multiple StSOR views (Fig. 4B). The distribution of assigned 3D angles indicated a preferred orientation but otherwise covered the entire 3D angular space (Fig. 4C). The cryo-EM analysis was conducted with octahedral symmetry from *ab initio* 3D reconstruction to the final 3D refinement and model reconstruction (see Materials and methods). With the final result, the gold-standard resolution with 0.143 FSC criterion was estimated to be ∼2.24 Å (Fig. 4D). The local 3D resolution ranged from ∼2.22 Å to ∼2.53 Å (Fig. 4E). The local resolution around the chimney structure was the lowest (approximately 2.4-2.5Å).

**Table 4.** Data collection, processing, and reconstruction statistics for cryo-EM.

| Data set | StSOR cryo-EM structure |
| --- | --- |
| **Data collection and processing** |  |
| Microscope | FEI Talos Arctica |
| Voltage (kV) | 200 |
| Detector | Falcon 3EC |
| Magnification | 150 k |
| Pixel size (Å) (calibrated) | 0.69 (0.676) |
| Electron dose (e<sup>-</sup>/Å<sup>2</sup>) | 1.00 |
| Number of frames | 50 |
| Defocus range | -0.2 to -1.5 |
| Number of collected micrographs | 2,558 |
| Number of selected micrographs | 2,387 |
| Number of particles for 2D classification | 305,182 |
| Number of particles for 3D classification | 176,437 |
| Number of particles for 3D refinement | 85,621 |
| Map Resolution (Å) | 2.24 |
| FSC threshold | 0.143 |
| Local resolution range | 2.22–2.53 |
| **Reconstruction** |  |
| Symmetry imposed | O |
| B factor (Å<sup>2</sup>) | 20.65 |
| Model composition |  |
| Protein | 60,120 |
| Ligand | 24 |
| Water | 2,243 |
| RMSD from ideal values |  |
| Bond length (Å) | 0.006 |
| Bond angle (°) | 0.834 |
| Ramachandran plot (%) |  |
| Favored/allowed/outlier | 96.75/3.25/0 |
| MolProbity score | 1.34 |
| Clashscore | 3.45 |
| Rotamer outliers (%) | 0 |
| CC<sub>mask</sub> | 0.89 |
| EMDB/PDB codes | EMD-30073/6M3X |

**Fig. 4.**
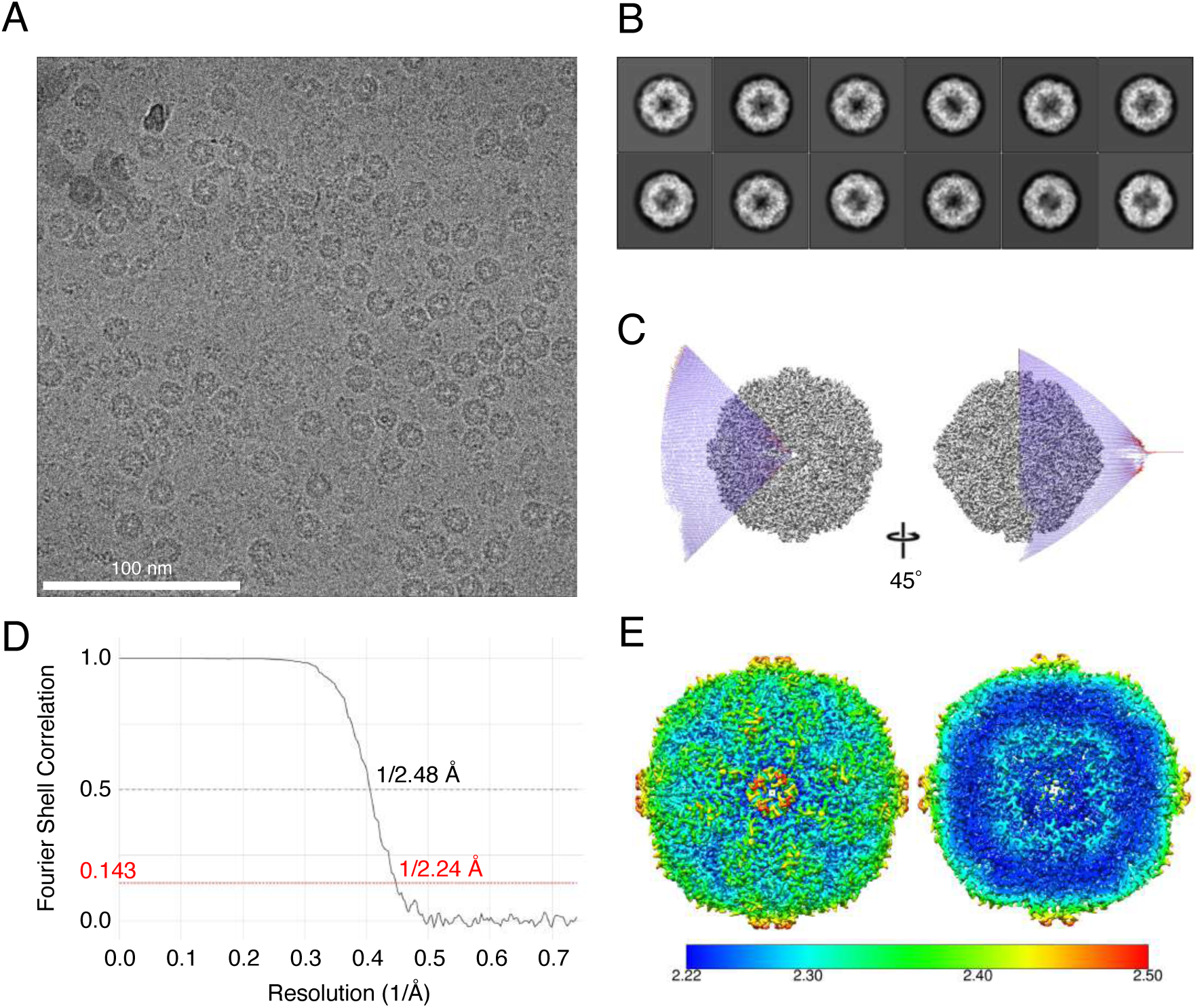
Single particle analysis of StSOR. (A) A representative micrograph of cryo-EM (scale bar 100 nm). (B) Representative two-dimensional class averages. (C) The sharpened density map of the final reconstruction with the resolution of 2.24Å (0.143 FSC criterion) and the angular distributions. (D) The Fourier Shell Correlation (FSC) curve. (E) The local resolution of the final structure.

The reconstructed cryo-EM structure consisted of 24 identical polypeptides with residues from Pro2 to Leu311. The MolProbity score, clashscore, and CC_mask_ (the correlation between the model and map) were 1.34, 3.45, and 0.89, respectively (Table 4). The overall cryo-EM structure of StSOR was almost the same as indicated by the crystal structure (Fig. 5A). The RMSD between the cryo-EM and crystal structures is 0.263 Å (for 309 Cα atoms). The RMSD values for the cryo-EM structure of StSOR in comparison with the crystal structures of AaSOR and AtSOR are ∼0.44 Å (for 260 Cα atoms) and ∼0.47 Å (for 267 Cα atoms), respectively. Fig. S5 shows the structure around three *cis*-peptides of this protein as examples of cryo-EM map.

**Fig. 5.**
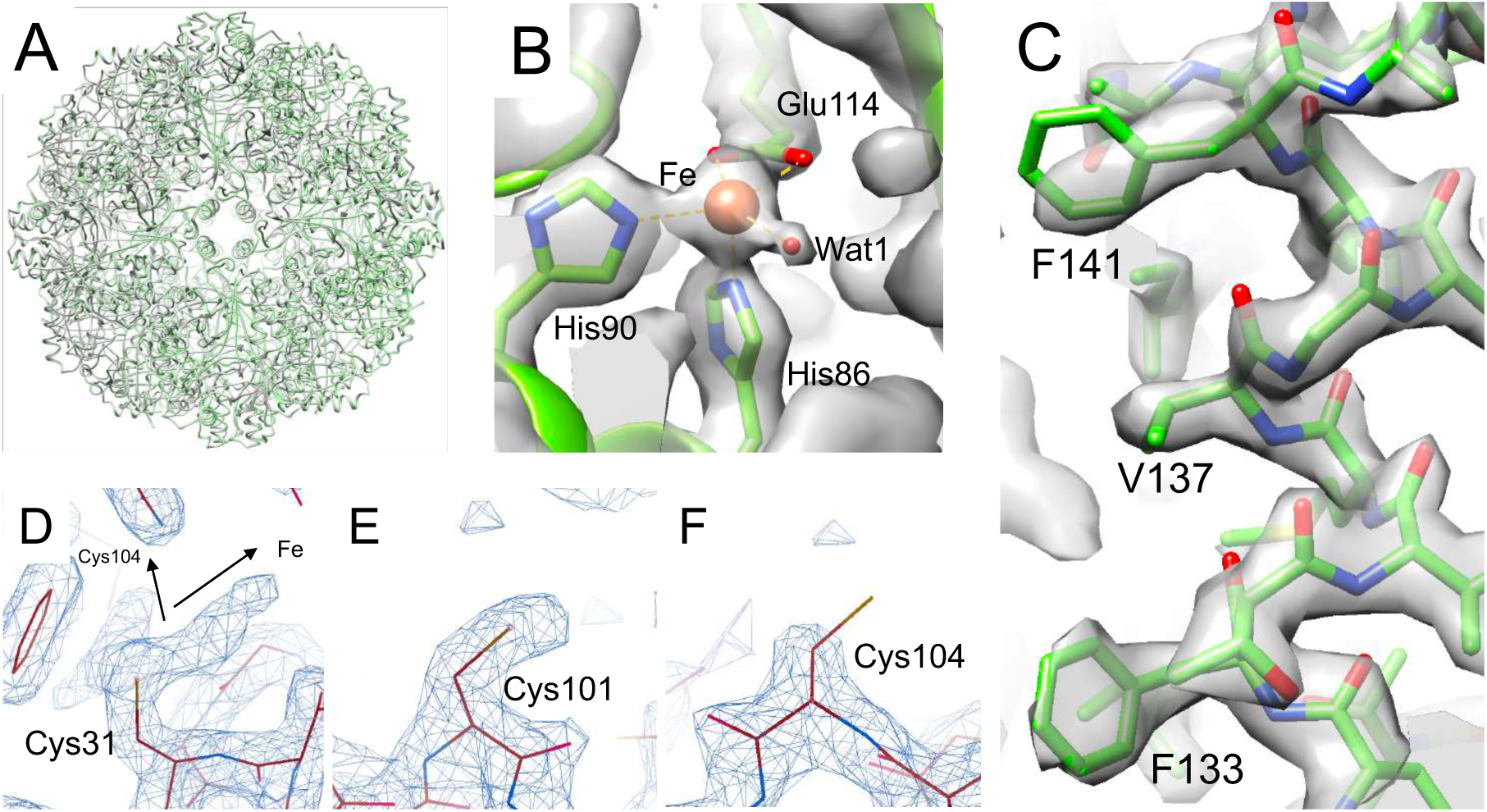
Cryo-EM structure of StSOR. (A) Superimposition of cryo-EM structure (green) and crystal structure (gray). RMSD was 0.263 Å for 309 Cα atoms. (B) Iron center with a contour level of 0.0435. (C) Residues forming a chimney with a contour level of 0.0435. (D-F) Cryo-EM map of the cysteines in the cavity. Maps of Cys31 (D), Cys101 (E), and Cys104 (F) are shown with a contour level of 0.0435 (4.5σ).

Protein models around the iron center fitted well to the cryo-EM map (Fig. 5B). The ligands were similarly located as those of the crystal structure, except that Wat2 was not observed. The CheckMyMetal server indicated trigonal bipyramidal 5-coordination with an outlier nVECSUM of 0.26 and the distances between the iron atom and the ligands were slightly greater than those in the crystal structure (Table 3 and Fig. S4B). In the chimneys, the main chain atoms of the helix fitted well to the map (Fig. 5C). However, their hydrophobic side chains were ambiguous. This observation may suggest that the side chains adopted various conformations for those particles vitrified in the cryo-EM grid, although some artifacts of cryo-EM analysis, e.g., errors in image alignments or damage caused by exposure to the liquid-gas interface are also possible. The trimer channels (discussed below) and other architectures showed no significant differences between the cryo-EM and crystal structures.

As shown in Fig. 5D-F, the cryo-EM maps around Cys31 and Cys104 exhibited different features compared with the crystal structure, while Cys101 was also observed as a non-flexible free thiol. The map around Cys31 is extended with a branch (Fig. 5D). One extension is oriented toward the iron center, while the other is oriented toward Cys104. The lengths of the extensions toward the iron and Cys104 are ∼4.0 Å and ∼3.4 Å, respectively at a 0.0435 (4.5σ) contour level. These lengths correspond to approximately two disulfide bonds (2.05 Å for each), thus implying formation of a persulfide, trisulfide, or branched oligosulfides. On the other hand, the Sγ atom of Cys104 lacks in the cryo-EM map for the same contour level (Fig. 5F), possibly because cysteine residues are prone to damage from electron beam exposures (Hattne et al., 2018).

## 4. Discussion

### 4.1 Substrate entry and exit pathways

The pathways for substrate entry and exit have been major topics of discussion regarding SOR structures because catalysis takes place inside the hollow chamber that is separated from the cytoplasmic environment (Li et al., 2008; Veith et al., 2011). See Fig. 2 in (Veith et al., 2011) for an overview of the substrate entry and exit pathways of SORs. In this section, we discuss the structural features of such pathways mainly by referencing the crystal structure of StSOR. The chimney is a major candidate for the entrance of S^0^ (Urich et al., 2006) and it is referred to as a “tetramer channel” (Li et al., 2008). Fig. 6A shows a surface model of four subunits forming the chimney. The size of the opening for this StSOR channel is similar to that of AaSOR. Phe133, Val137, and Phe141 are lined inside the chimney to form a hydrophobic channel that is suitable for S^0^ entry (Fig. 6B). An electron density map shows that the hydrophobic residues are clearly observed in the crystal structure (Fig. 6C) and they are basically conserved in SORs from various organisms (Fig. 1B). Other channels formed by three subunits that are arranged along a 3-fold axis, referred to as “trimer channels”, were suggested to be exit routes for the reaction products (HSO_3_^−^, HS^−^ and S_2_O_3_^2-^) because of their polar characteristics in AtSOR and AaSOR (Li et al., 2008). The diameters of the trimer channels in StSOR are (Fig. 6D) comparable to those of the chimneys, and three hydrophilic residues (Arg99, Gln103, and Asn225) are involved in the trimer channel. The residues involved in the trimer channel are not conserved except for Arg99 (Fig. 1B). In particular, Met227 contributes to the formation of a more hydrophobic and narrower channel compared with AaSOR and AtSOR (Fig. 6E and 6F). For the active site pore that connects the active site pocket and the central cavity of the chamber of the 24-mer (Fig. 3D), alanine substitution of the two conserved key methionine residues (Met296 and Met297, see Fig. 1B) in AaSOR significantly lowered the activity (Veith et al., 2011). Two histidine residues (His166 and His277) to which Zn^2+^ binds and cause an inhibitory effect on AaSOR (Veith et al., 2011) are also conserved in StSOR (Fig. 1B). There were no significant structural differences at the Zn^2+^ binding site of these SORs. However, Zn^2+^ did not inhibit the StSOR activities (Fig. S2A).

**Fig. 6.**
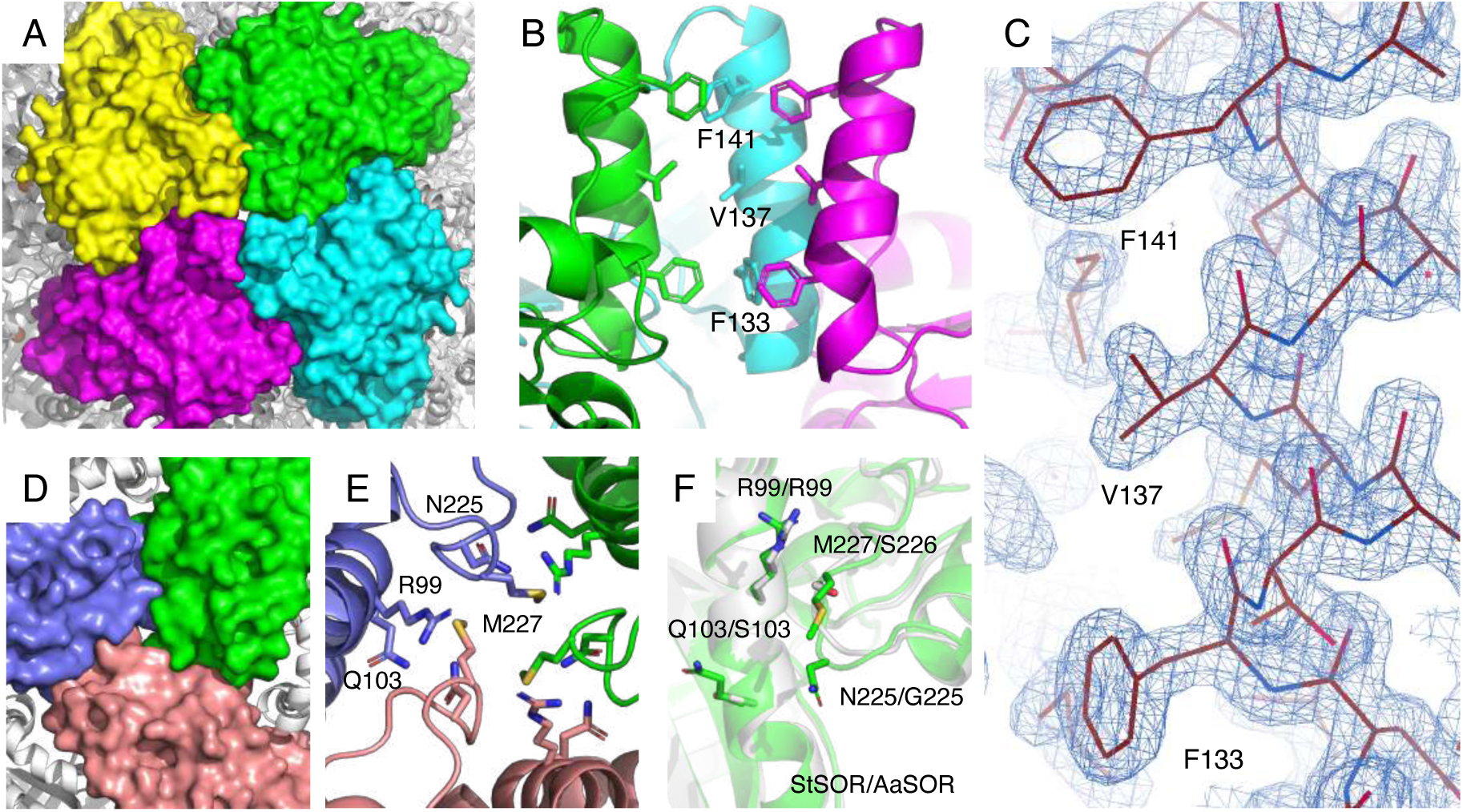
Channels to the cavity (chamber) of the crystal structure of StSOR. (A) Molecular surface model of four subunits forming the chimney (tetramer channel). (B) Hydrophobic residues lined inside the tetramer channel. For clarity, one subunit is not shown. (C) Electron density map of residues forming the chimney with a contour level of 1.3σ. (D) Molecular surface model of three subunits forming the trimer channel. (E) Residues forming the trimer channel. (F) Superimposition of StSOR (green) and AaSOR (grey). Side chains of the residues forming the trimer channel are shown as sticks.

### 4.2 Comparison of the crystallographic and cryo-EM structures

Structural analyses of StSOR by two distinct methodologies (X-ray crystallography and cryo-EM) enabled us to compare the three-dimensional structures with and without crystal packing effects. The hydrophobic side chains of the chimneys were clearly observed in the electron density map of the crystal structure (Fig. 6C). Due to the crystal packing effect (Fig. S6A), the chimneys have relatively low B-factor values compared to the interior regions of the hollow sphere (Fig. S6BC). On the other hand, chimneys were determined with lower local resolution than inside the molecule in the cryo-EM structure (Fig. 4E), and the map of the side chains are more ambiguous (Fig. 5C). This feature may reflect the functionally flexible nature of the chimneys as entrances of the rather bulky polysulfide substrate(s) (Urich et al., 2006).

The StSOR structures as determined by two structural methodologies highlight the differences in some catalytic components. In the cryo-EM structure (single particle analysis), one of the two iron-ligating waters (Wat2) was not observed. Considering the crystal structure of AtSOR in which the Wat1 site is not occupied (Li et al., 2008), the coordination spheres around the iron may change during the SOR catalytic cycle. Further structural analysis of other SORs will clarify the various ligation states of iron in relation to the catalytic cycle. In addition, Cys31 is a critical residue for catalytic reactions by StSOR (Table 1). As with the case of AtSOR (Li et al., 2008), Cys31 in the crystal structure of StSOR does not have any additional electron densities. On the other hand, a branched extra map (blob) next to the side chain of Cys31 exists in the cryo-EM structure (Fig. 5D). The map does not resemble the one observed in the AaSOR crystal structure, which was interpreted as cysteine persulfide (Urich et al., 2006). We cannot suggest any plausible assignments for the map but a subtle difference of chemical conditions in crystallization and preparation of a grid or effects of photoreduction and electron beam exposure during the data collections might influence Cys31 sensitivity. Cys104 is located at a relatively distant position (13 Å) from the iron center (Fig. 3E) but its activity is significantly reduced by the alanine substitution (Table 1). Cys104 may act as a substantial support for enzymatic functions related to side chain flexibility that are assumed from the present cryo-EM structure (Fig. 5F). Li et al. postulated that Cys101 and Cys104 are involved in sulfur disproportionation reactions (Li et al., 2008). However, this hypothesis is not supported by our mutational study because the decreases in oxygenase and reductase activities of C101A and C104A mutants were nearly parallel (Table 1). Although no mechanistic roles of Cys101 and Cys104 have been suggested, these residues may support catalysis by considering the form of substrate (S^0^), such as linear polysulfides, that can extend across the iron and cysteines (Urich et al., 2006). Otherwise, participation in the entry, orientation, and exit of inorganic sulfur substances is intended.

### 4.3 StSOR as a benchmark protein of cryo-EM measurements

In this report, we determined the X-ray crystal structure and the cryo-EM structure of StSOR at 1.73 Å and 2.24 Å resolutions, respectively (Tables 2 and 4). Although the meaning of “resolution” of the two different methods is not the same, the coordinates and isotropic temperature factors of the protein atoms were determined with sufficient quality for analyzing the functional components of the enzyme. In particular, the resolution of the cryo-EM structure was fine enough to determine the side chain conformation (Fig. 5) and *cis*-peptide bonds (Fig. S5). Previously, Herzik et al. reviewed examples of single-particle cryo-EM structures which were resolved to better than 3 Å using a 200 kV transmission electron microscope and argued their cost-effectiveness compared to 300 kV instruments (Herzik et al., 2017). Here, we compared the resolutions of cryo-EM structures determined by single-particle analysis using 200 kV instruments, which are currently available on EMDB (Table S1). The resolution of our StSOR structure is one the highest for cryo-EM structures and resolutions better than 2.2 Å were achieved only using mouse heavy-chain apoferritin (Danev et al., 2019) and rabbit aldolase. Considering the high thermostability (Fig. 2A) and molecular symmetry, the good visibility of the partially purified sample in the micrographs (Figs. S1B and 4A) and the angular distributions of the particles covering all orientations (Fig. 4C), StSOR can be used as a benchmark sample for EM facilities as an alternative to the widely used apoferritin.

## Abbreviations

SOR: sulfur oxygenase reductase;
AaSOR: *Acidianus ambivalens* SOR;
AqSOR: *Aquifex aeolicus* SOR;
AtSOR: *Acidianus tengchongensis* SOR;
cryo-EM: cryogenic electron microscopy;
CTF: contrast transfer function;
DTNB: 5,5’-dithiobis(2-nitrobenzoic acid);
FSC: Fourier shell correlation;
HnSOR: *Halothiobacillus neapolitanus* SOR;
PAGE: polyacrylamide gel electrophoresis;
*p*CMB: *p*-chloromercuribenzoate;
RMSD: root mean square deviation;
SbSOR: *Sulfobacillus thermosulfidooxidans* SOR;
SD: standard deviation;
SDS: sodium dodecyl sulfate;
StSOR: *Sulfurisphaera tokodaii* SOR;
TpSOR: *Thioalkalivibrio paradoxus* SOR.

## Accession numbers

The EM map of StSOR has been deposited to the Electron Microscopy Data Bank (EMDB: EMD-30073). The coordinates of the cryo-EM structure have been deposited to the Protein Data Bank (PDB: 6M3X). The atomic coordinates and structure factors of the StSOR crystal structure have been deposited (PDB: 6M35).

## Acknowledgments

The authors thank the staff of the Photon Factory for the X-ray data collection, Prof. Mitsuru Abo for ICPAES measurements, and Chiho Masuda and Rieko Sukegawa for helpful assistance during the cryo-EM study. This study was supported by JSPS-KAKENHI (24580136 to T. W.) and the Platform Project for Supporting Drug Discovery and Life Science Research [Basis for Supporting Innovative Drug Discovery and Life Science Research (BINDS)] from AMED under Grant Number JP20am0101071 (support number 1400 and 2114). This work was partially supported by JSPS-KAKENHI (19H00929, 15H02443, 26660083, and 24380053 to S. F.).

