## Supplementary Methods, Tables, and Figures for "Crystallographic and cryogenic electron microscopic structures and enzymatic characterization of sulfur oxygenase reductase from *Sulfurisphaera tokodaii*"

**<u>Contains:</u>**

- **Supplementary Table S1**
- **Supplementary Figures S1-S6**
- **Supplementary Methods**
- **References**

**Table S1.** High resolution single-particle cryo-EM structures in the databases with data collected by 200 kV instruments.

| Resolution (Å) | Sample | Source organism | Oligomeric state | Theoretical weight (MDa) | EMDB ID | PDB ID | Citation |
|---|---|---|---|---|---|---|---|
| 1.75 | Apoferritin | Mouse | 24 | 0.505 | 21024 | 6v21 | bioRxiv 855643 |
| 2.01 | Apoferritin | Mouse | 24 | 0.44 | 9914 | none | (Danev et al., 2019) |
| 2.13 | Aldolase | Rabbit | 4 | 0.15 | 21023 | 6v20 | bioRxiv 855643 |
| 2.24 | StSOR | *S. tokodaii* | 24 | 0.55 | 30073 | 6m3x | This work |
| 2.32 | HemQ | *Geobacillus* sp. | 5 | 0.144 | 21373 | 6vsa | bioRxiv 798280 |
| 2.56 | Capsid protein VP1 | Adeno-assisted virus | 60 | 5.08 | 20693 | 6u95 | (Kaelber et al., 2020) |

Prepared based on database searches on 21 February 2020.

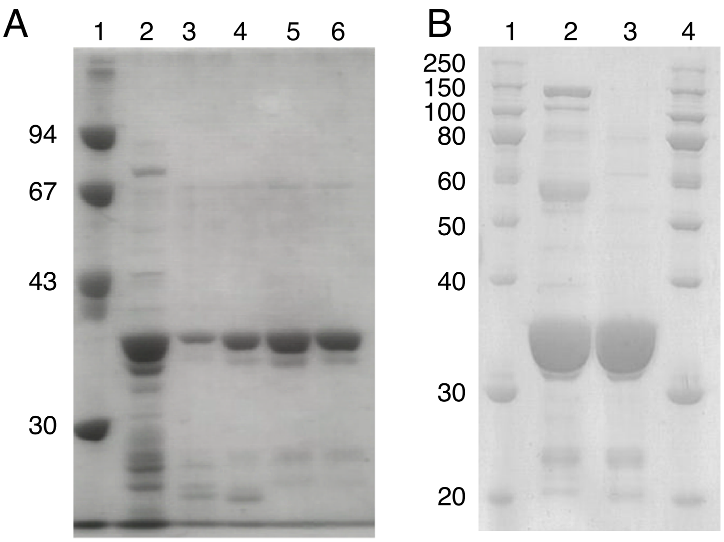


**Figure S1.** SDS-PAGE of purified recombinant samples. SDS-PAGE of purified protein samples for biochemical (A) and structural (B) characterizations are shown. (A) 1, marker; 2, crude cell extract; 3, after Q-Sepharose; 4, after Hydroxyapatite; 5, after Mono Q; 6, after Superdex 200pg. (B) 1 and 4, marker; 2, after Superdex 200pg that was used for cryo-EM; 3, after Mono Q that was used for crystallization.

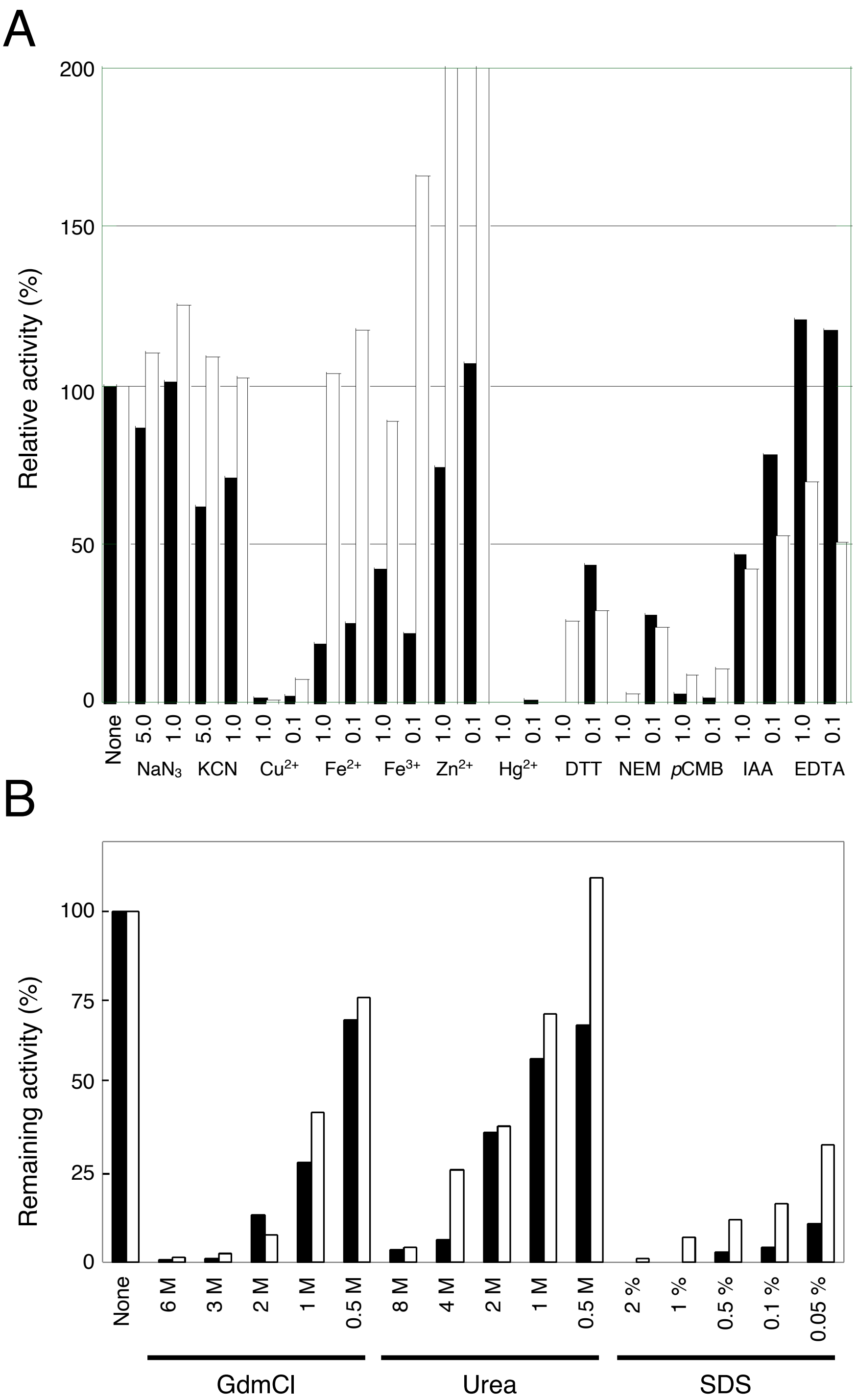


**Figure S2.** Effects of inhibitors, metals, and denaturants. (A) Effect of inhibitors and metal ions. The activities in the presence of indicated concentrations (mM) of each additive were measured. Metal ions were added as chloride salts. Abbreviations: DTT, dithiothreitol; GdmCl, guanidium chloride; NEM, *N*-ethylmaleimide; *p*CMB, *p*-chloromercuribenzoate; IAA, iodoacetic acid; EDTA, ethylenediaminetetraacetic acid. Relative activities in the presence of 0.1 and 1.0 mM $Zn^{2+}$ were 318% and 252%, respectively. (B) Stability against denaturants. Remaining activities after incubation for 30 min at 4 °C in the presence of each denaturants were measured. Symbols: filled bars, oxygenase activity; open bars, reductase activity.

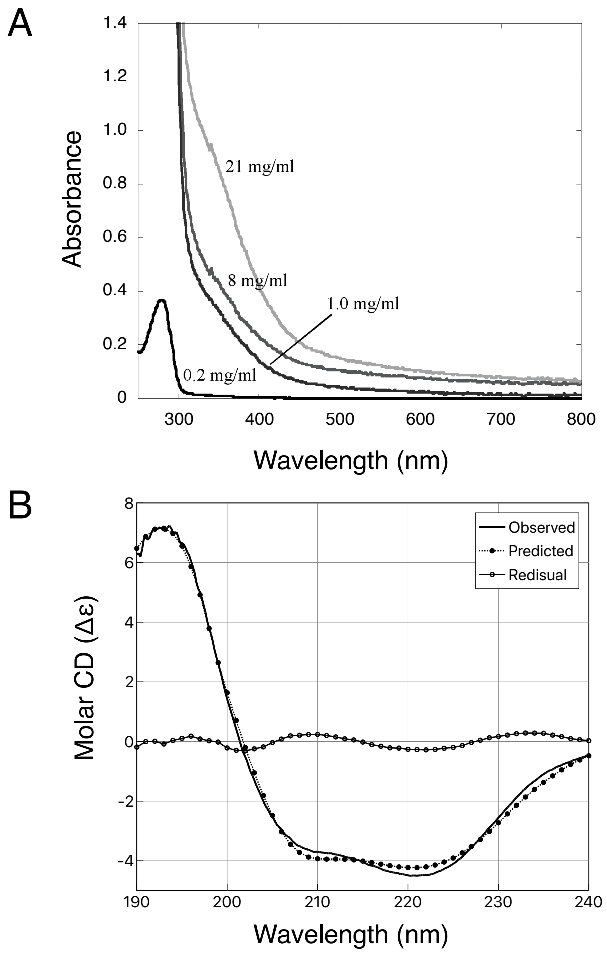


**Figure S3.** Spectral analyses of purified StSOR. (A) UV-visible absorption spectra in 10 mM Tris-HCl (pH 8.0) at room temperature (25 °C). Protein concentrations are indicated. (B) CD spectrum of 0.25 mg/mL StSOR in 10 mM Na-phosphate buffer (pH 7.0) at room temperature.

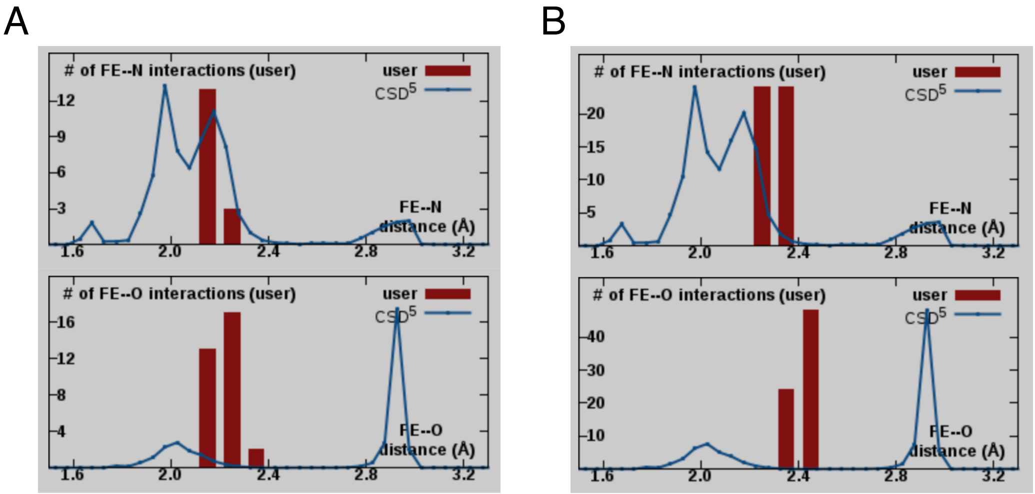


**Figure S4.** Metal (Fe)-ligand (N or O) distance distribution of crystallographic (A) and cyro-EM (B) structures analyzed by CheckMyMetal server. Distance distributions in the StSOR structures (user, red bars) and Cambridge Structural Database (CSD, blue lines) are shown.

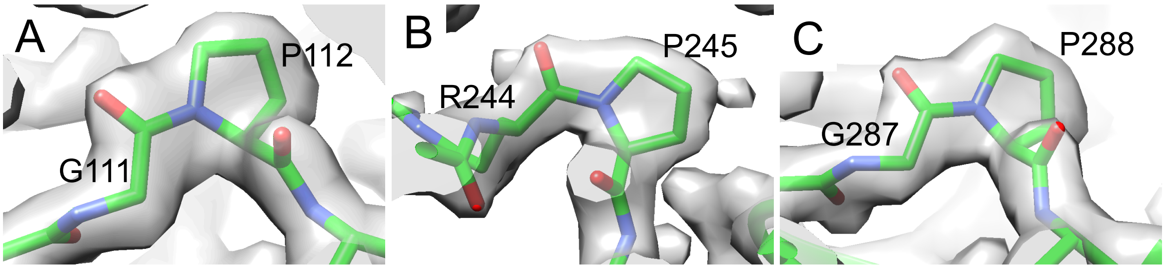


**Figure S5.** Cryo-EM structure of cis-peptides. Density maps of Gly111–Pro112 (A), Arg244–Pro245 (B), and Gly287–Pro288 (C) are shown with a contour level of 0.0435 (4.5σ).

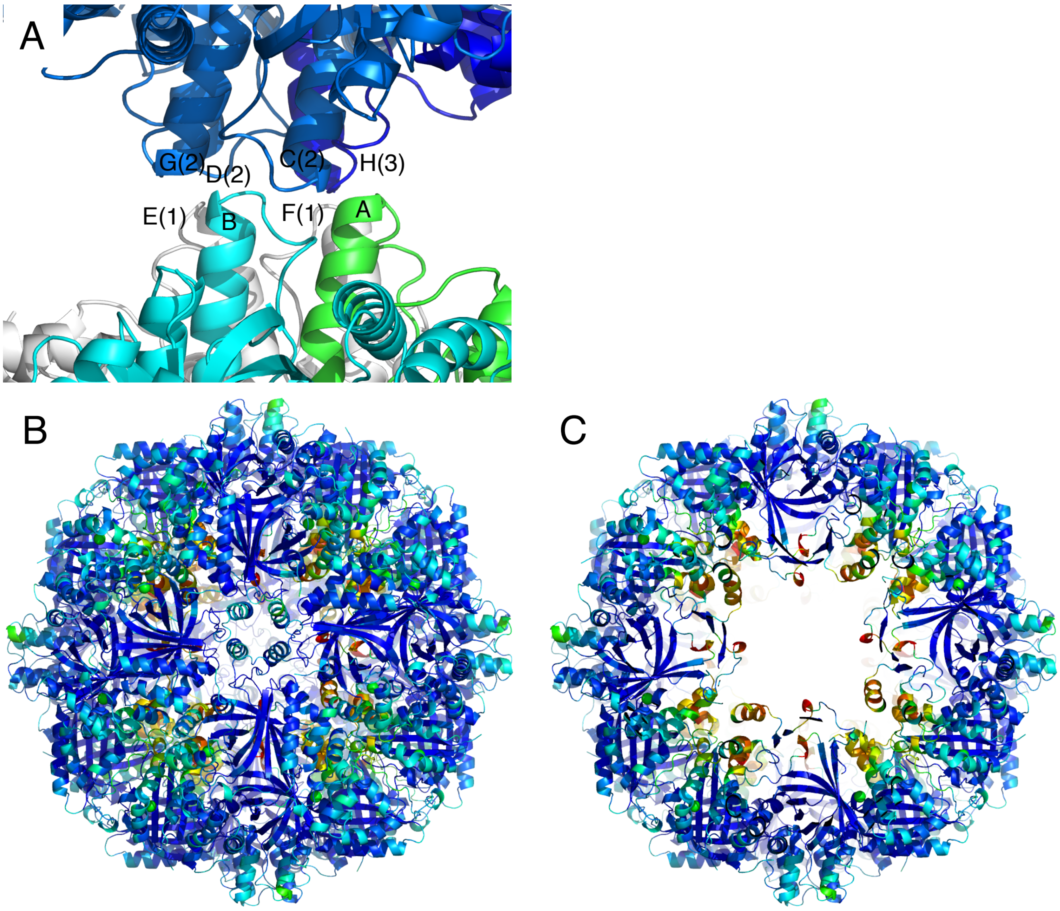


**Figure S6.** Packing effects and temperature factors of the crystal structure. (A) Packing interaction of chimneys. Chain A (green) and B (cyan) of the original model coordinates and symmetry-related molecules are shown. Chains E and F in symmetry molecule 1, chains C, D, and G in molecule 2, and chain H in molecule 3 are shown in white, marine, and blue, respectively. (B and C) The 24-mer structure of StSOR colored by B-factor as a rainbow color from blue (20 Å$^2$) to red (60 Å$^2$). Panel (B) shows the molecular surface with no clipping, and panel (C) shows inside of the hollow sphere by clipping with 80 Å slab.

### Supplementary Methods

#### Cryo-EM data processing

The movie frames were aligned, dose-weighted, and averaged using Motioncor2 (Zheng et al., 2017), on 5 × 5 tiled frames with a B-factor of 200 applied, to correct for beam-induced specimen motion and to account for radiation damage by applying an exposure-dependent filter. The micrographs whose total accumulated motion was larger than 100 Å were discarded. The non-weighted movie sums were used for Contrast Transfer Function (CTF) estimation (512-pixel box size, 30 Å minimum resolution, 3 Å maximum resolution, 0.10 amplitude contrast) with CTFFIND4 (Rohou and Grigorieff, 2015) and Gctf program (Zhang, 2016), while the dose-weighted sums were used for all subsequent steps of image processing. The images whose CTF max resolution was better than 5.5 Å were selected. In addition, micrographs showing obvious indication of ice crystallization (i.e. strong ice ring in the Fourier space) were manually discarded.

The particles were picked fully-automatically using SPHIRE-crYOLO with the generalized model (Moriya et al., 2017; Wagner et al., 2019) with 231-pixel box size and the selection threshold of 0.05. The micrographs that contain less than 36 picks were excluded. A stack of 305,182 particle images was extracted from 2,390 dose-weighted sum micrographs while rescaling to 2.76 Å/pixel with 96-pixel box size, and subjected to consecutive two runs of reference-free 2D classification (1$^{st}$ run: 200 expected classes, 188 Å mask diameter; 2$^{nd}$ run: 200 expected classes, 166 Å mask diameter) using RELION-3 (Zivanov et al., 2018). The 160,571 particles corresponding to the best 29 classes of 2$^{nd}$ run that displayed secondary-structural elements and multiple views of StSOR were selected for RELION-3 *ab initio* reconstruction (asymmetry, single expected class, 188 Å mask diameter), and 176,437 particles are selected with more relaxed criteria for the subsequent RELION-3 3D classification (2 expected classes). The generated *ab initio* map was imposed octahedral symmetry, low-pass filtered to 20 Å, and used as an initial model for the 3D classification. The 3D class obviously consisted from bad images containing non-targeted objects and was removed, and the volume of the best 3D class was low-pass filtered to 15 Å, and used as an initial model for the subsequent RELION-3 3D refinement (octahedral symmetry, 240 Å mask diameter, with padding) with the 152,484 selected particles.

Then, the refined volume was rescaled to 0.69 Å/pixel with 400-pixel box size, low-pass filtered to 15 Å, and used for the subsequent 3D refinement. Accordingly, the selected particle images were also re-centered and re-extracted using the same rescale settings, and 3D auto-refined (octahedral symmetry, 240 Å mask diameter, no padding) twice, the 1st run without 3D mask and the 2nd with a soft-edged 3D mask created from the initial 3D reference of 1st run (5-pixel extension, 10-pixel soft cosine edge). The 2nd run resulted in the resolution of 2.88 Å. 152,457 particles were again re-centered and re-extracted without changing the rescale settings. To refine per-particle defocus, beam tilt, and beam-induced motion corrections, the cycle of CTF refinement and Bayesian polishing (Zivanov et al., 2019) in RELION-3 was repeated four times. To measure degree of the improvement, 3D refinement (octahedral symmetry, 240 Å mask diameter, no padding) with the volume of previous run as initial 3D reference, a soft-edged 3D mask created from the initial reference (5-pixel extension, 10-pixel soft cosine edge) and solvent-flattened FSCs options was used after each CTF refinement and Bayesian polishing step. The 3D refinement after 4th Bayesian polishing run yielded the resolution of 2.44 Å.

At this point, the pixel size was checked by comparing the 2.44 Å cryo-EM structure with the atomic-coordinate model (PDB code: 6M35) using "Fit in Map" tool in UCFS Chimera (Pettersen et al., 2004), and calibrated to 0.676 Å/pixel which minimized the number of "atoms outside contour". Using the calibrated pixel size, 146,465 selected particles were again re-centered and re-extracted with 480-pixel box size from 2,387 dose-weighted sum micrographs. Accordingly, the density map was also rescaled with the same settings, low-pass filtered to 15 Å, and used as an initial model for the subsequent 3D refinement (octahedral symmetry, 236 Å mask diameter, no padding) with a soft-edged 3D mask (5-pixel extension, 10-pixel soft cosine edge). Then, the two cycles of CTF refinement and Bayesian polishing steps were executed and generated the 2.25 Å structure.

To improve homogeneity of the particle stack, no-alignment 3D classification was conducted by setting expected classes to 2 and regularization parameter T to 16 (octahedral symmetry, 236 Å mask diameter, no padding) with a soft-edged 3D mask (5-pixel extension, 10-pixel soft cosine edge), and selected 85,621 particles by choosing the 3D class with the best resolution. The last 3D refinement (octahedral symmetry, 236 Å mask diameter, no padding) with a soft-edged 3D mask (5-pixel extension, 10-pixel soft cosine edge) and solvent-flattened FSCs option generated the final result of 2.24 Å resolution. The re-runs of the last Bayesian polishing step excluding various numbers of the last movie frames to adjust the total electron dose did not improve the reconstruction. The local resolution of the final reconstruction was estimated using the RELION-3's own implementation of ResMap (Kucukelbir et al., 2014).

For calculation of the global resolution estimation after each 3D refinement, the gold-standard FSC resolution with 0.143 criterion (Rosenthal and Henderson, 2003) was used, including the phase randomization to account for the possible artifactual resolution enhancement caused by solvent mask (Chen et al., 2013). For the visualization of the output 2D/3D images, UCFS Chimera and e2display.py of EMAN2 (Tang et al., 2007) were used. To calculate the smallest box size that ensures no CTF aliasing in the reciprocal space up to an expected resolution for the maximum defocus value of the dataset, the ctflimit function (Penczek et al., 2014) implemented in SPARX/SPHIRE (Hohn et al., 2007; Moriya et al., 2017) was used.
